# Spatial organization, grouping strategies and cyclic dominance in asymmetric predator-prey games

**DOI:** 10.1101/090316

**Authors:** Annette Cazaubiel, Alessandra F. Lütz, Jeferson J. Arenzon

**Author notes:** Electronic address (Annette Cazaubiel). Electronic address (Alessandra F. Lütz). Electronic address (Jeferson J. Arenzon).

## Abstract

Predators may attack isolated or grouped preys in a cooperative, collective way. Whether a gregarious behavior is advantageous to each species depends on several conditions and game theory is a useful tool to deal with such a problem. We here extend the Lett-Auger-Gaillard model [Theor. Pop. Biol. **65**, 263 (2004)] to spatially distributed groups and compare the resulting behavior with their mean field predictions for the coevolving densities of predator and prey strategies. We show that the coexistence phase in which both strategies for each group are present is stable because of an effective, cyclic dominance behavior similar to a well studied generalizations of the Rock-Paper-Scissors game with four species (without neutral pairs), a further example of how ubiquitous this mechanism is. In addition, inside the coexistence phase (but interestingly, only for finite size systems) there is a realization of the survival of the weakest effect that is triggered by a percolation crossover.

## I. INTRODUCTION

There is a myriad of foraging strategies that predators utilize to increase their rate of success. Among them, preys may be attacked in a cooperative, coordinated way by a group of predators, whose similar actions are correlated in space and time. When this involves diffierent and complementary behaviors, it is also called a collaboration [1]. Examples of such coordinated or collaborative hunting are lions [2–4] (including the pair of man-eaters lions of Tsavo [5]), hawks [6], crocodiles [7], spiders [8, 9], and several other species [1]. Interspecies collaborations exist as well (for example, between fishermen and dolphins in the south of Brazil [10, 11] or between honey-hunters men and honeyguide birds in Mozambique [12], among others [13]). Hunting in group may bring several benefits and has been widely discussed (for a review, see Ref. [1] and references therein). For example, it increases the probability of capturing a large prey [6, 14, 15], helps prevent the carcass to be stolen by other predators [16, 17], allows for faster spotting [18] and more complex distracting, tracking and chasing tactics, helps related conspecific that may be unable to hunt or are in the process of acquiring hunting skills [15, 19], etc. On the other hand, there may be costs as it also increases the competition between members of the group while feeding, concentrates the search for food to a smaller territory what may decrease the number of available preys, etc. Grouping tactics may also benefit preys [20]. Surveillance is more efficient when done in parallel by several individuals while the others have more time to feed themselves [21–23]. The probability of being caught is smaller [24, 25] and the group may take advantage of group distracting [26], intimidating and escaping techniques. On the other hand, a group of preys may be more easily spotted than an isolated one and the resources should be shared by all members [27, 28]. In addition to those factors, for both preys and predators, collective decision making can be improved in larger groups [29, 30] (but information sharing may involve costs [31] and benefits [32] as well).

Despite those mounting experimental results, much less attention has been dedicated to model coordinate hunting [33]. Recently, Lett et al [34] (hereafter referred as the LAG model) introduced a game theory model in which the abundances of preys and predators were assumed constant and only the fractions of those populations using either an individual or collective strategy coevolved (see, however, Ref. [35]). The LAG model takes into account some of the advantages and disadvantages for both preys and predators choosing a grouping strategy. More specifically, it is assumed that grouping lowers the risk of being preyed at the cost of increasing the competition for resources, while predators have a greater probability of success at the expense of having to share the prey with others, sometimes referred to as the “many-eyes, many mouths” trade off [27, 28]. Preys and predators were modeled by assuming fully mixing (no spatial structure), a mean field approach, and the temporal evolution of both densities being described by replicator equations [36].

A complementary approach, based on a less coarse grained description, explicitly considers the spatial distribution of individuals and groups. The local interactions between them introduce correlations that may translate into spatial organization favoring either grouping or isolated strategies, raising a number of questions. For instance, do these strategies coexist within predators or preys populations? If yes, is this coexistence asymptotically stable? How does the existence of a local group induce or prevent grouping behavior on neighboring individuals? Do gregarious individuals segregate, forming extended regions dominated by groups? In other words, how spatially heterogeneous is the system? Does the replicator equation provide a good description for both the dynamics and the asymptotic state? If not, when does it fail? If many strategies persist, which is the underlying mechanism that sustains the coexistence? We try to answer some of these questions with a version of the LAG model in which space is explicitly taken into account through a square lattice whose sites represent a small sub-population patch. Each of the sites is large enough to contain only a single group of predators and preys at the same time. If any of these groups is ever disrupted, their members will resort to a solitary strategy, hunting or defending themselves alone.

The paper is organized as follows. We first review, in Section II A, the LAG model [34] and summarize the main results obtained with the replicators equation, and then describe, Section II B, the agent based implementation with local competition. The results obtained in this diffierent framework are presented in Sec. III. Finally, we discuss our conclusions in Section IV.

## II. THE MODEL

### A. Replicator Equations

Lett et al [34] considered, within a game theoretical framework, grouping strategies for preys and predators. Both can choose between single and collective behavior and each choice involves either gains or losses for the individuals, as discussed in the introduction. The relevant parameters of the model are defined in Table I. The size of both populations is kept constant during the evolution of the system, only the proportion of cooperative predators, *x*(*t*), and the fraction of gregarious preys, *y*(*t*), evolve in time (see, however, Ref. [35] for a version that also considers population dynamics). Whether they increase or decrease depends on how their payoff compare with the average payoff of the respective whole population. If collective behavior earns a larger payoff than the average, the associated density increases, otherwise it decreases. This dynamics is then described by the replicator equations [36].

**Table I:** Model parameters [34] along with the default value considered here.

| Symb. | Definition and default value |
| --- | --- |
| $p$ | probab. of a group capturing a lone prey (0.5) |
| $G$ | gain per captured prey per unit of time (1) |
| $n$ | predators in a group (3) |
| $e$ | preys captured by a group of predators (2) |
| $\alpha$ | preying efficiency loss due to grouped preys |
| $\beta$ | preying efficiency loss when hunting alone |
| $\gamma$ | reduction of prey resources due to aggregation (1) |
| $F$ | gain for isolated preys per unit of time (1) |

For the fraction *x* of predators hunting collectively, the payoff is [34]

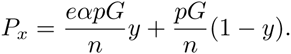

The first contribution comes from the interaction of these predators with the fraction *y* of preys that organize into groups for defense. By better defending themselves, the preys reduce the hunting efficiency by a factor *α*, nonetheless *e* preys are captured with probability *p* and the gain *G* per prey is shared among the *n* members of the group of predators. The second term is the gain when the group attacks an isolated prey, whose density is 1*−y*, and shares it among the *n* predators as well. When the remaining 1*−x* predators solely hunt, they are limited to a single prey and an efficiency that is further reduced by a factor *β*, what is somehow compensated by not having to share it with others. This information is summarized in the payoff matrix:

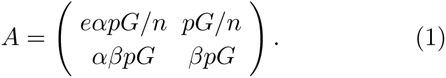

As isolated preys consume the available resources, the gain per unit of time, on average, is *F*. Once aggregated, the resources are shared and the individual gain reduced by a factor *γ*. The fraction of preys that aggregates becomes less prone to be preyed by a factor *α*. If the grouped preys are attacked by a group of predators, *e* preys are captured and Lett et al [34] considered that the payoff coefficient is 1 − *eαp* (imposing *eαp* ≤ 1). On the other hand, a lone predator has its efficiency reduced by a factor *β*, thus the surviving probability is 1 − *βp* or 1 − *αβp* for an individual or a group of preys, respectively. The payoff for the fraction *y* of preys that remain grouped is then written as

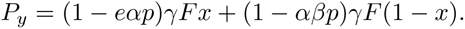

A similar consideration can be done for isolated preys and the payoff matrix for preys is

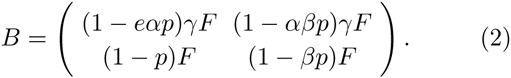

It is the difference between the payoff *P* and its average, *P¯*, that drives the evolution of both *x* and *y*. Indeed, the replicator equations, 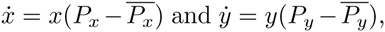 giving the rate at which these two densities evolve in time, are [34]

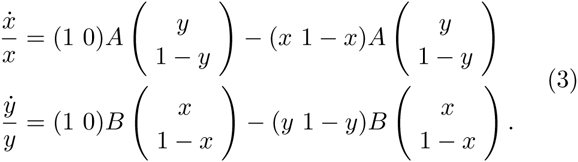

These equations describe an asymmetric game and can be rewritten as [36]:

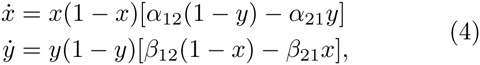

where

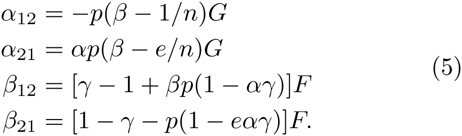

Eqs. (4) have five fixed points: the vertices of the unit square and the coexistence state (*x*\*, *y*\*):

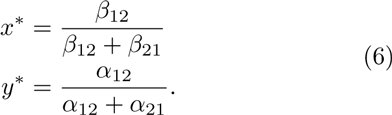

The asymptotic state is determined only by the signs of *α*_12_, *α*_21_, *β*_12_ and *β*_21_ [36]. Indeed, if *α*_12_*α*_21_ < 0 or *β*_12_*β*_21_ < 0, the densities of grouped predators *x* and grouped preys *y* will monotonously converge to an absorbent state in which at least one of the populations is grouped, i.e, (0,1), (1,0) or (1,1). We use the notation (*x*_*∞*_, *y*_*∞*_) for the asymptotic state (e.g, *x*_*∞*_ ≡ *x*(*t → ∞*)) of the whole system and *x*_*∞*_ *y*_*∞*_ for the site variable describing the combined state of site *i*. On the other hand, if *α*_12_*β*_21_ > 0 and *β*_12_*β*_21_ > 0, there are two diffierent possibilities. From the linear stability analysis [34, 36], (*x*\*, *y*\*) is a saddle point if *α*_12_*β*_12_ > 0, and the system ends up in one of the vertices of the unit square. On the other hand, if *α*_12_*β*_12_ < 0, the eigenvalues are imaginary and the system evolves along closed orbits around the center (*x*\*, *y*\*). In other words, when this last condition is obeyed, both strategies, grouped or not, coexist at all times with oscillating fractions of the population and the time average of such behavior, once in the stationary regime, corresponds to the point Eq. (6).

The above replicator equations predict that both *x* and *y* approach the asymptotic state 1 or 0 exponentially fast (except in the coexistence phase). For instance, as discussed below, inside phase (1,1), *y* attains the asymptotic state much faster than *x*. Then, taking *y* = 1 and expanding for small 1 − *x*, we get *x*(*t*) ∝ 1 − exp(−*α*_21_*t*). The characteristic time is 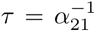 and as *β* → *e/n*, *τ* diverges as *τ ∼* (*βn* − *e*)^−1^. Diffierent transitions may depend on other coefficients, Eq. (5), but the exponent at the transition is always 1.

Besides presenting general results, Lett et al [34] also discussed the particular case when there is no reduction in resource intake by the preys when they are grouped (*γ* = 1). In this case, their behavior is a response to the capture rate alone. By also considering *F* = *G* = 1, we see that while *β*_12_ is always positive, *β*_21_, *α*_12_ and *α*_21_ change sign at *eα* = 1, *βn* = 1 and *βn* = *e*, respectively. These changes in sign lead to diffierent asymptotic be-haviors (phases) and locate the transition lines between them. As expected, preys are grouped when *α* is small, whatever the value of *β*. Similarly, small values of *β* lead to cooperating predators for all values of *α*. Remarkably, for *eα* > 1 and intermediate values 1 < *βn* < *e*, the eigenvalues of the Jacobian associated with Eqs. (4) become purely imaginary [34]. This corresponds to a coexistence phase where the densities of grouped animals oscillate in time along closed orbits around the center point (*x*\*, *y*\*) given by Eq. (6), as discussed above.

### B. Spatially Distributed Population

The above description of the competition between collective and individual strategies for both predators and preys does not take into account possible spatial correlations and geometrical effects. Space is usually introduced by considering an agent based model in which individuals are placed on a lattice or distributed on a continuous region. The unit cell corresponds to the smallest viable group and on each site of the lattice there may exist one or none of such groups. Since both predators and preys coexist in each site, there are two variables (*x*_*i*_, *y*_*i*_), *i* = 1, …, *N* (where *N* = *L*^2^ is the total number of sites and *L* is the linear length of the square lattice) that takes only the values 1 or 0, the former when agents are grouped, the latter for independent individuals.

Diffierently from the mean-field description where pay-off was gained from interactions with all individuals in the system, alone or grouped, in the lattice version interactions are local and occur only between nearest neighbors sites (self-interaction is also considered since each site has both predators and preys). At each step of the simulation, one site (*i*) and one of its neighbors (*j*) are randomly chosen. The predators (preys) on *i* interact with the preys (predators) both on *i* and in all neighboring sites, accumulating the payoff 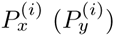. At the same time, both groups in *j* also accumulate their payoffs. The updating involves the site with the smallest payoff adopting the strategy of the other with a probability proportional to the difference of payoffs. For example, for predators (and analogously for preys), if *P_x_^(j)^* > *P_x_^(i)^*, the probability that *i* changes its state is

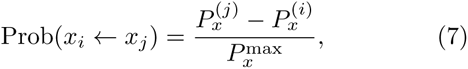

where 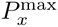 is the maximum value of the accumulated payoff of the predators for the chosen parameters. This rule is known to recover the replicator equation when going from the microscopic, agent based scale to the macroscopic, coarse grained level [37].

## III. RESULTS

Fig. 1 shows the temporal evolution of both *x*(*t*) and *y*(*t*) for *α* = 0.2, several values of 0 < *β* < 1 and an initial state with *x*(0) = *y*(0) = 1/2. In this case, for all values of *β*, preys remain mostly grouped at all times since *y*(*t*) > 0.5 (bottom panel). As predicted by mean field, there is a transition at *β* = 2/3 where predators change strategy (top panel): for *β* > 2/3, hunting alone becomes efficient and does not involve sharing the prey, thus *x*_*∞*_ = 0. On the other hand, when the cost of sharing the prey is compensated by a more efficient preying, *β* < 2/3, we have *x*_*∞*_ = 1. Interestingly, for the initial state chosen here, the behavior of *x*(*t*) is not monotonous when (1 + 2*α*)/3(1 + *α*) < *β* < 2/3: *x*(*t*) initially decreases, *x*(0) < 0, until attaining a minimum value and then resumes the increase towards *x*_*∞*_ = 1. The location of this minimum corresponds to the time at which *y* crosses the point *y*\*, and the envelope of all minima goes along with the plateau developed for *β* > 2/3 as this value is approached from above. In this latter region, the behavior follows a two steps curve: there is a first, fast approach to the plateau followed by the departure from it at a much longer timescale. In this model, the fast relaxations occur as preys organize themselves into groups (increasing *y*) while the later slow relaxation is a property of the predators alone and is caused by the orbit passing nearby an unstable fixed point, as can be understood from Eqs. (4). For values of in the interval (1+2*α*)/3(1+*α*) < *β* < 2/3, if *y* < *y*\*, *x* decreases until *y* crosses the line at *y*\*, whose value depends on both and *β*. At this point, the coefficient of ˙*x* is zero and there is a minimum. Once *y* > *y*\*, *x* resumes the increase and approaches the asymptotic state *x*_*∞*_ = 1 exponentially fast. As *β* → 2/3^−^, *y*\* → 1 and the minimum crosses over to an inflection point. Indeed, for *β* > 2/3, *x*(*t*) decreases towards 0 after it crosses the plateau. This behavior is seen both in the simulations and in mean field.

**FIG. 1:**
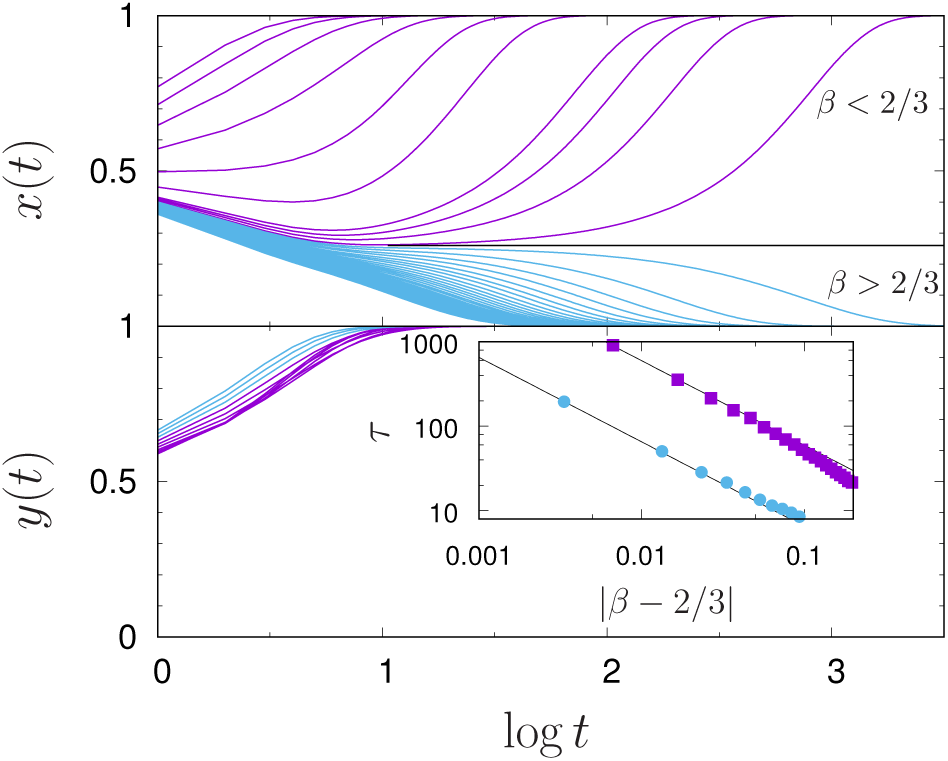
Behavior of *x*(*t*) and *y*(*t*) as a function of time for *α* = 0.2, *L* = 100 and several values of *β*. From both mean-field and simulations, the transition from collectively hunting predators to single, individual hunt occurs at *β* = *e/n* = 2/3. Notice that most of the preys are in the aggregated state at all times, with *y*(*t*) monotonously increasing from *y*(0) = 0.5 to *y*(*1*) = 1. Predators, on the other hand, have a richer behavior (see text). As = 2/3 is approached from both sides, a plateau at *x* ≃ 0.26 develops (the dashed, horizontal line is only a guide to the eyes). Inset: Power-law behavior of the characteristic time *τ* such that |*x* − *x*_∞_| < 0.2 around the transition at *β* = 2/3. The straight lines have exponent 1, as predicted by the replicator equation, although the coefficients di er by one order of magnitude. The top curve is for *β* → 2/3^−^ while the bottom one is for *β* → 2/3^+^.

Associated with the late exponential regime there is a characteristic time that diverges as a phase transition is approached. For example, in mean field, 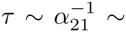 (*βn* − *e*)^−1^ for the (1,1)-(0,1) transition. In the simulations, as *x* approaches the limiting value *x*_*∞*_, *τ* is estimated as the time beyond which |*x* − *x*_∞_| < *ε*, where 0 < *ε* < 1 is chosen, for convenience, to be *ε* = 0.2. The exponent measured in the simulations is in agreement with the mean field prediction, as can be seen in the inset of Fig. 1.

For *α* = 0.8 there is, in agreement with the replicator equation, a third phase for 1/3 < *β* < 2/3 (see Fig. 2). Diffierently from the previous case, in this intermediate region, both strategies may persistently coexist. There is an initial, transient regime in which both *x* and *y* oscillate and, depending on and *β*, the orbit may get very close to one or both absorbing states (0 and 1), i.e. heteroclinic. Because of the stochastic nature of the finite system, sometimes it ends in one of those absorbing states during the oscillating regime, otherwise the amplitude of the oscillations decreases and a mixed fixed point is eventually attained. A natural question is how close this fixed point is from the mean field prediction, Eq. (3). Since it is the coexistence phase that presents new, non trivial behavior, we discuss it in detail now.

**FIG. 2:**
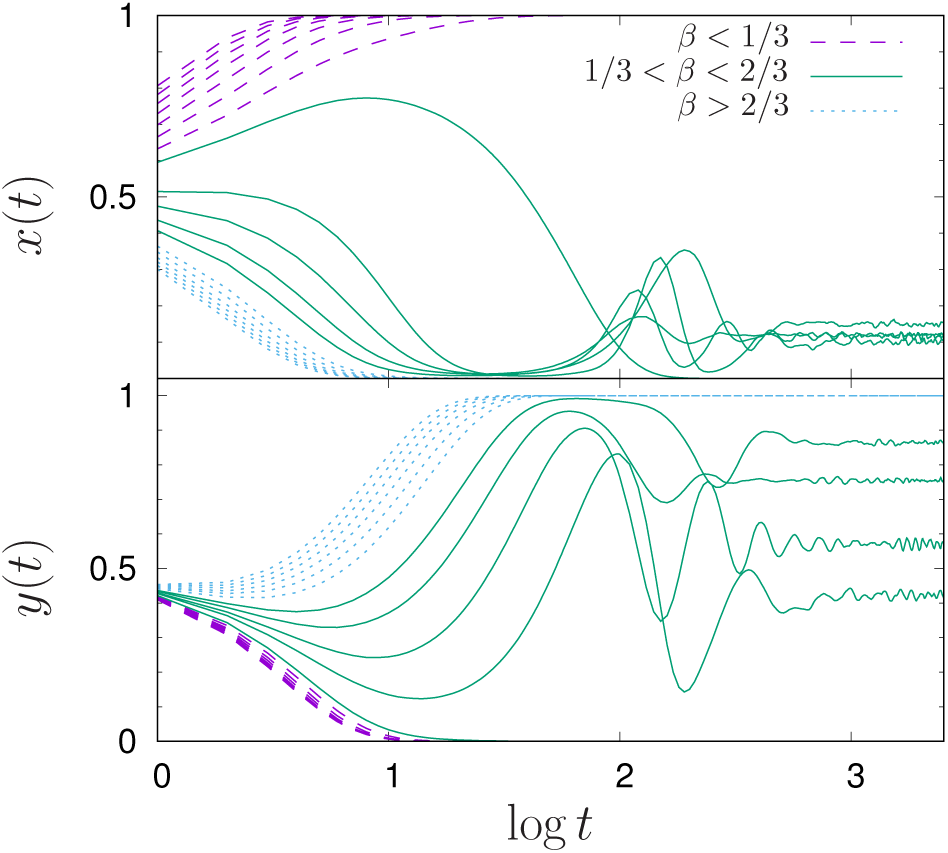
The same as in Fig. 1 but for = 0.8. Two transitions are present at *β* = 1/*n* = 1/3 and *e/n* = 2/3, with a coexistence phase in between.

The mean field behavior can be observed in the solid lines of Fig. 3 as the system enters the coexistence region at a fixed *β* = 0.4 from the (1,1) phase. This scenario completely changes when predator and preys are spatially distributed. While mean field predicts that *x*_*∞*_ and *y*_*∞*_ present a continuous and discontinuous transition, respectively, at 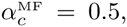 the simulation shows continuous transitions for both quantities at a smaller value, *α*_*c*_ ≃ 0.45. This is not a finite size effect since the curves, for the system sizes considered here, collapse onto a single curve in this region up to *α* ≃ 0.6 (and the larger the system is, the wider the collapsed region becomes). Both solid curves monotonically decrease with α in the coexistence phase: while *x*_*∞*_ smoothly goes from 1 down to 0, *y*_*∞*_ barely changes and the curve is almost flat (e.g., for *β* = 0.4, 1/5 < *y*_∞_ < 1/3). Not only there is little correspondence between this predicted behavior for (*x*_*∞*_, *y*_*∞*_), but even the overall, qualitative behavior is diffierent. In the spatial model, *y*_*∞*_ does not present a monotonous, having instead a minimum at *α* ≃ 0.6, even in the limit of very large systems. The existence of this minimum is remarkable because one would expect that as predators become more efficient against grouped preys, the fraction of such preys would decrease, exactly what mean field predicts. Both *x*_*∞*_ and *y*_*∞*_ present strong finite size effects in this phase. During the time evolution whose asymptotic state is shown in Fig. 3, above a size dependent value of *α*, *x* is absorbed onto the group disrupted state (*x*_*∞*_ = 0) and, immediately after, *y*_*∞*_ evolves toward 1. The interval in which these effects occur (i.e. where the (0,1) phase for *β* > 2/3 is reentrant inside the coexistence phase) decreases with *N* and seems to vanish in the thermodynamical limit (we will get back to this point below). Assuming that the trend shown in Fig. 3 continues as the system approaches the thermodynamic (deterministic) limit, one can build a phase diagram as shown in Fig. 4. For example, the (0,1) phase that, for finite systems, extends below the line *β* = 2/3, seems to be suppressed in the thermodynamic limit. The only transition line that seems to disagree with the mean field is the reentrant coexistence phase around *α* = 1/2 and 1/3 < *β* < 1/2, that is not, as mentioned above, a finite size effect. No (0, 0) phase exists neither in the simulation nor in mean field.

**FIG. 3:**
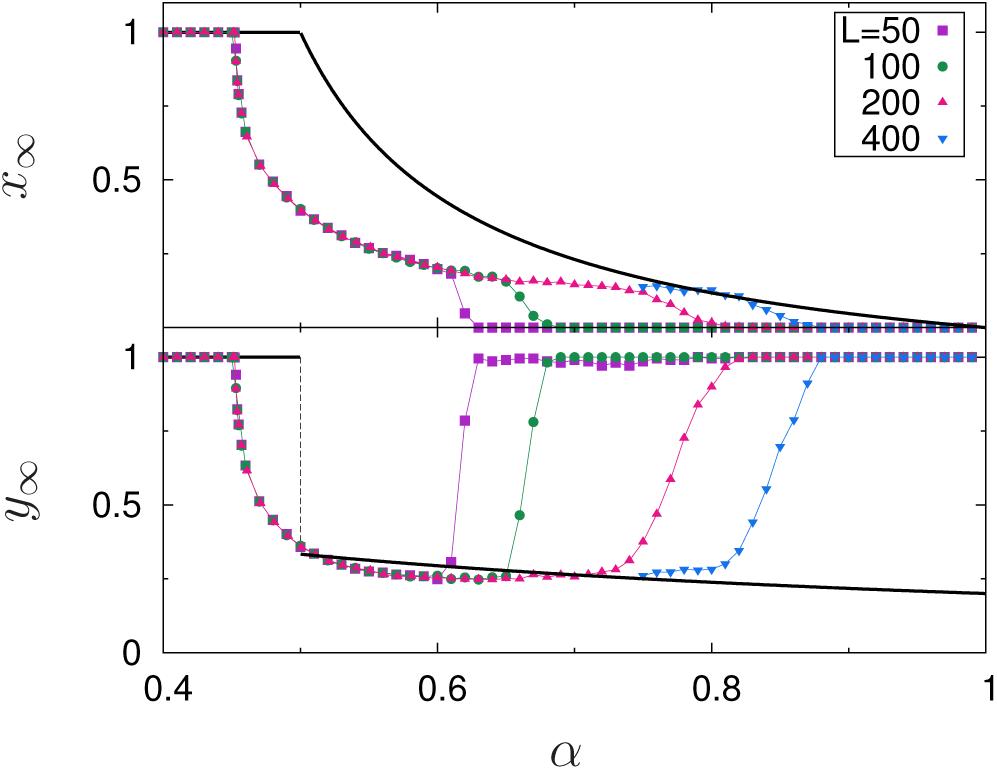
Asymptotic value of *x*(*t*) (top) and *y*(*t*) (bottom), for *β* = 0.4, as a function of *α*. The solid lines correspond to the fixed points, Eqs. (6), predicted in mean field and discussed in Sect. II A. Strong finite size effects occur for large values of where the system is absorbed onto the (0,1) state. Both transitions are continuous and occur at a smaller *α*_*c*_ than predicted in mean field.

**FIG. 4:**
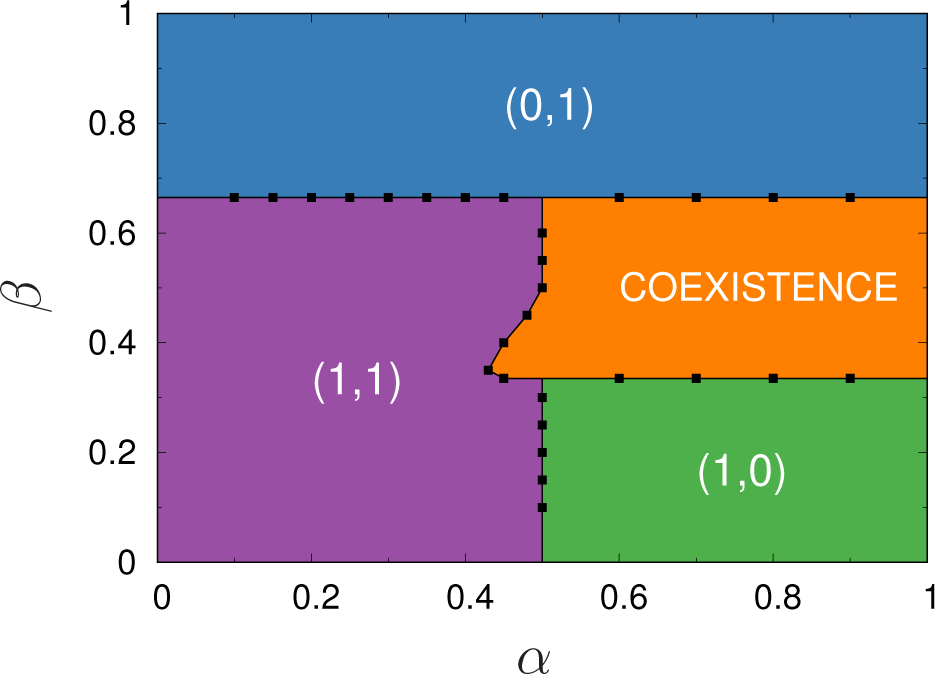
Phase diagram showing the diffierent phases for the spatial version after extrapolating the boundaries to the deterministic limit. The reentrant coexistence phase is at odds with the replicator equations that predicts a vertical line at *α* = 1/2 for 0 ≤ *β* ≤ 2/3. The horizontal lines, on the other hand, do agree with mean-field. Notice that only the (0,0) fixed point does not appear and is replaced by the coexistence phase.

In the coexistence phase, any of the four combinations of lone and collective strategies for both predators and preys may be present at each site (*x*_*i*_ *y*_*i*_): 00, 01, 10 and 11. Fig. 5 shows, for = 0.4 and diffierent values of *α*, how these four strategies form domains whose sizes become larger with *α*. While the coexistence persists, the 00 strategy forms the percolating background against which the other three struggle to survive by spatially organizing themselves in compact, nested domains whose borders move, invading other domains in a cyclic way. In Fig. 6a we show the direction of these invasions by placing, in the initial state (*t* = 0, top row), one strategy inside, and other outside a circular patch. The bottom row shows the corresponding state after 40 Monte Carlo steps. By switching positions, the invasion direction is reversed, indicating that it is not a simple curvature driven dynamics but, instead, involves a domination relation. Taking into account the six combinations of Fig. 6a, the interaction (or flow) graph shown in Fig. 6b summarizes how the diffierent strategies interact (this particular orientation of the arrows may change for other points in the coexistence phase [38]). Similar cyclic dominance behavior has been extensively studied in predator-prey models with interaction graphs with three (Rock-Paper-Scissors) or more species [37, 39] with intransitive (sub-)loops. Indeed, the topology of the interaction (or flow) graph in Fig. 6b is closely related to the one in Ref. [40]. The invasion is either direct as in the first four columns (or, equivalently, along the perimeter of the interaction graph) or, as seen in the last two columns, involves the creation of an intermediate domain. For example, a patch of 00s, does invade 11 (fifth column in Fig. 6a) by first disrupting the preys organization, thus, the strategy 10 appears, invades 11 and, in turn, is invaded by 00. In those cases that the invasion proceeds through an intermediate domain (that is diffierent from the diagonal strategies being neutral), we use a dashed arrow in the interaction graph. This cyclic dominance among the combined strategies of preys and predators is the underlying mechanism that explains the persistence of the coexistence state in this region of the phase diagram.

**FIG. 5:**
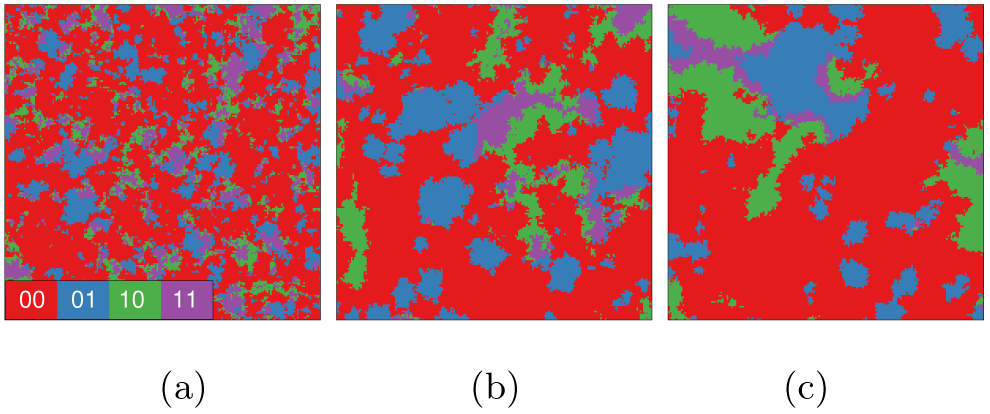
Snapshots using a color code for the combined *x*_*i*_ *y*_*i*_ strategy for *β* = 0.4 and *α* = 0.6 (left), 0.7 (middle) and 0.8 (right), inside the coexistence phase. Notice that the strategies 10-01-11 organize into large intertwined domains, whose characteristic size increases with *α*, against a 00 background.

Another characteristic of the coexistence phase is the presence of strong finite size effects, as shown in Fig. 3 and discussed above. Again, this behavior can be explained observing the patterns in Fig. 5. Upon the background of lonely strategies (00) there are combined clusters of the 01+10+11 strategies whose characteristic size increases with (from left to right in the figure). Fig. 5c was prepared for = 0.8, just below the percolation threshold where, for *L* = 400, the 00 cluster no longer percolates (while the combination of the other three strategies does). Interestingly, for the system sizes considered in Fig. 3, the size dependent percolation threshold coincides [38] with the value of where *x*_*∞*_ drops to 0 and *y*_*∞*_ attains 1. That is, once the combined cluster of 01+10+11 strategies percolates, disrupting the background 00 cluster, 01 is the only surviving strategy. This is a percolation triggered, finite size realization of the survival of the weakest effect [41]. The sudden decrease of the 00 population effectively reduces the 01 invasion rate what, as the name says, turns it into the only surviving strategy. Diffierently from the model in Ref. [40], here the crossed interactions are not direct, as discussed above. Thus, even if 01 has two predators and a single prey (00), this prey also helps by eliminating 11, that also happens to prey on 01. Thus, these favorable conditions for 01 contribute for its persistence. When the characteristic linear size of the combined cluster 01+10+11, that increases with *α*, becomes larger than *L*, it percolates and the system evolves until it is absorbed onto the state 01. By further increasing *L*, it takes a larger to achieve the percolation threshold, explaining the strong finite size effects observed. It is important to emphasize that the invasion graph in Fig. 6 was obtained for the particular values *β* = 0.4 and *α* = 0.7. Other points inside the coexistence phase may show similar behavior albeit with some arrows reversed and strategies switching roles while keeping some of the sub-loops intransitive and the coexistence stable [38].

**FIG. 6:**
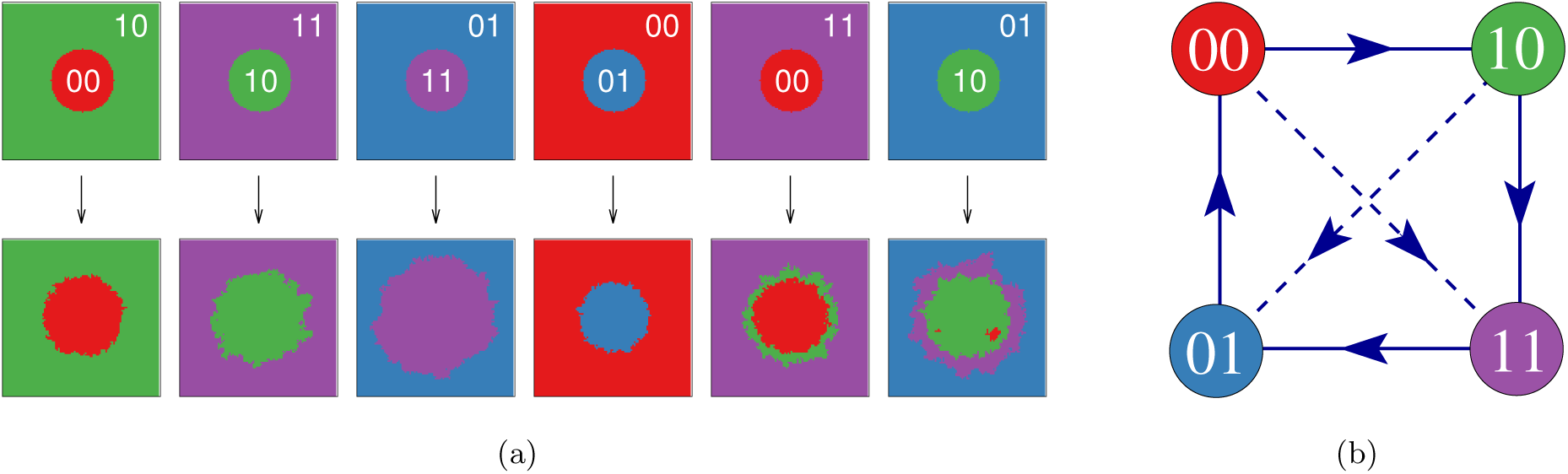
a) Dominance of strategies for = 0.4 and = 0.7 (inside the coexistence phase) starting from an initial, circular patch (top row) we show the state after 40 MCS. In some cases, denoted by the solid arrow in the invasion graph at the right, the initial patch increases in size. For the interactions along the diagonal (dashed arrows), the invasion occurs in two steps. For example (fifth column), 00 first disrupts the aggregated preys, forming an intermediate 10 cluster, and then grows into it. Notice that for the 10 invading 01 (last column), besides the strategy 11 that intermediates the invasion, there are some sources of 00 growing inside the 10 patch (at the interface 01-10, all combinations may be created and, sometimes, migrate toward the interior of the circle). b) Interaction (flow) graph showing the direction of the invasion front for each of the four strategies.

## IV. CONCLUSIONS AND DISCUSSION

The foraging behavior of predators and the corresponding defensive response from preys is fundamental to understand how small animal communities spatially distribute, organize and eventually allow for more complex forms of sociability [42–44]. It thus becomes important, when studying simple models for such behavior, to go beyond mean field where a fully mixed, infinite size system is considered and the spatial structure, with the local correlations it implies, is missing. We considered here a finite dimensional stochastic version of the model introduced by Lett et al [34] in which short range interactions between predators and preys are taken into account. In this way, spatial correlations that change, to some extent, the foraging behavior predicted by the replicator (mean field) equation, and the accompanying new dynamical behavior are introduced. The game theoretical framework for this model considers two strategies, collective or individual, for both hunting predators and defensive preys. The advantages and disadvantages of each option are modeled by a set of independent parameters from which we considered only two: how group preying efficiency is reduced against grouped preys (*α*) and the reduction factor of lone predators (*β*). We then study in detail this particular case, also discussed within mean field in Ref. [34], in the limit of high population viscosity in which strategy dispersion is a slow process, solely occurring due to the newborn limited dispersal driven by the pairwise updating rule between nearest neighbors patches.

The extensive simulation results presented here confirm that the overall phase diagram is essentially the same both in mean field and in finite dimension (but infinite size). The combined strategies of both predators and preys present four possible states in our binary version. There are three phases in which either preys or predators (or both) behave collectively, (1,0), (0,1) and (1,1), while the (0,0), albeit also absorbing, is not prevalent in any region of the phase diagram. Instead, a coexistence phase appears with 00 sites forming a percolating background upon which compact clusters of the other strategies distribute and compete (for the particular and used here). More importantly, our results unveil the underlying mechanism not only for the strategies coexistence in this phase, but also for the occurring strong finite size effects. Specifically, this model is an example of an asymmetric game presenting cyclic dominance, between the above four combined strategies, in the coexistence phase. Starting, say, with a 10 population of collective predators preying on lone individuals, free riders have the advantage of not having to share their preys with the other members of the group and the 00 strategy invades 10. At this point, if preys organize into groups they may better defend themselves, with 01 thus replacing 00. The lone predators feel the urge to prey collectively and 11 dominates over 01. Finally, 10 invades 11 because it is better for predators to go after grouped preys, since the number of captured preys (*e*) compensates for the loss of efficiency (*α*). Thus, is better for aggregated preys to stay alone. For = 0.7 and = 0.4, the strategies 11, 01, 11 form spatially extended domains whose borders are not static and move accordingly with the flow graph of Fig. 6b. This metacluster is embedded in a sea of 00 strategies (that are essential for coexistence because the 01-10-11 loop is transitive) up to a size dependent value of where it percolates and the subsequent evolution leads to the extinction of three strategies, the surviving one being the 01 state. Interestingly, since this effect is triggered by the initial reduction of the 00 strategy, the only prey of 01, when the metacluster percolates, this effective reduction of the preying rate of 01 is the responsible for its survival, a well known effect in cyclic games, the survival of the weakest [41].

Besides studying simple models for this less explored foraging variability, our results emphasize how important it is to study both the asymptotic states and the dynamics towards them in a finite size population. It is not possible to disentangle the strong finite size effects observed in the asymptotic state of the stochastic model from the dynamical evolution since it is the very existence of heteroclinic orbits that make the system prone to be captured by an absorbing state because of fluctuations. Since actual populations are far from the thermodynamical limit, the results obtained in the mesoscopic limit become relevant. Although preys rapidly converge to the asymptotic strategy, predators timescales may be very large and the transient coexistence may extend to times much larger than those that are relevant in practice. Moreover, the very existence of differences between the mean field and the spatial version shows the importance of trying several complementary approaches even in the study of quite simplified models. Because of the importance of the geometry of domains, it is interesting to examine their properties in more detail [38].

Preys and predators usually engage in somewhat coordinated chase and escape interactions [45–47] that also allow them to explore and profit from neighboring patches. It is thus interesting to check whether, and to what extent, the properties of the model, in particular in the coexistence region, do change in the presence of mobile individuals. Chasing and escaping behaviors may be quite complex depending on the physical and cognitive attributes of each individual and involve space and time correlations between the displacements and change in velocity of each agent [48]. Simple movement rules, if not completely random, are likely to generate repeatable (and, because of that, exploitable) patterns of behavior, while those involving higher levels of variability and complexity somehow involve more advanced cognitive skills. By studying such mobility patterns, one could get a better understanding of how important the cognitive abilities are in defining hunting strategies [1]. Moreover, these patterns are also important for the demographic distribution of both predators and preys since the shu ing of strategies, depending on how random [49] the mobility is, may decrease the spatial correlation and destroy local structures, changing the spatial organization of preys and predators [43].

Several other extensions of this model are possible. For simplicity, we assumed that the size of a group is constant and homogeneous throughout the population. However, it can also be considered as a dynamical parameter coevolving along the collectivist trait. Also, it would be interesting to further explore the possibilities offered by the spatial setup, for example, having heterogeneous parameters depending on the landscape (resources may not be evenly distributed) or due to the variability of the species. Another important question is how the size of the hunters group respond to an increase or stochastic fluctuations in the size of a swarm of preys (and vice-versa). Although we focus here on the binary situation, it is also possible to have larger patches with enough individuals to form more than one group, such that the variables describing the local populations may now become continuous. In addition, the present model does not consider noise, a simple, effective way of taking into account some of the missing characteristics that may influence the outcome. It would thus be interesting to see how robust the results are in the presence of noise. The short-range interactions present in the model are not able to synchronize distinct regions in the coexistence phase. By introducing a fraction of long-range connections we expect global oscillations to be restored, similarly to those found in mean-field [50, 51]. What is the threshold fraction of such interactions for having global oscillations and how it depends on the parameters of the model, along with the other points above, are still open questions.

## Acknowledgments

AC thanks the IF-UFRGS for the hospitality and the ENS-Paris for partial support during her stay in Porto Alegre. AFL is partially supported by a CNPq PhD grant. JJA thanks the INCT Sistemas Complexos and the Brazilian agencies CNPq, Fapergs and CAPES for partial support.

